# Predicting causal citations without full text

**DOI:** 10.1101/2022.07.05.498860

**Authors:** Travis A. Hoppe, Salsabil Arabi, B. Ian Hutchins

## Abstract

Insights from biomedical citation networks can be used to identify promising avenues for accelerating research and its downstream bench-to-bedside translation. Citation analysis generally assumes that each citation documents causal knowledge transfer that informed the conception, design, or execution of the main experiments. Citations may exist for other reasons. In this paper we identify a subset of citations that are unlikely to represent causal knowledge flow. Using a large, comprehensive feature set of open access data, we train a predictive model to identify such citations. The model relies only on the title, abstract, and reference set and not the full-text or future citations patterns, making it suitable for publications as soon as they are released, or those behind a paywall (the vast majority). We find that the model identifies, with high prediction scores, citations that were likely added during the peer review process, and conversely identifies with low prediction scores citations that are known to represent causal knowledge transfer. Using the model, we find that federally funded biomedical research publications represent 30% of the estimated causal knowledge transfer from basic studies to clinical research, even though these comprise only 10% of the literature, a three-fold overrepresentation in this important type of knowledge transfer. This finding underscores the importance of federal funding as a policy lever to improve human health.

**Significance statement:** Citation networks document knowledge flow across the literature, and insights from these networks are increasingly used to form science policy decisions. However, many citations are known to be not causally related to the inception, design, and execution of the citing study. This adds noise to the insights derived from these networks. Here, we show that it is possible to train a machine learning model to identify such citations, and that the model learns to identify known causal citations as well. We use this model to show that government funding drives a disproportionate amount of causal knowledge transfer from basic to clinical research. This result highlights a straightforward policy lever for accelerating improvements to human health: federal funding.

## Introduction

United States (US) federal science funders, research institutions, and investigators are jointly charged with advancing the frontiers of knowledge to stimulate innovation, improve human health, and maintain a competitive national edge in the global knowledge economy (1). In recent years, the generation of large open datasets of grants, publications, clinical trials, and patents have facilitated the generation of linked knowledge networks that relate resources to knowledge creation and eventual flow into applied outcomes (2–5). Insights from these networks, about the most effective approaches in achieving these important research goals, can be used to identify promising avenues for accelerating research and its downstream applied impacts (6). However, despite the power of analyzing large-scale knowledge flow to inform the decision-making of scientists, institutions and funders, these efforts are complicated by one simple fact: many, if not most, of citations in the scientific literature represent transfer of information that did not directly influence the inception, design, or execution of a research study. This makes it more challenging than would otherwise be the case to identify the subset of citations in the literature that really drove subsequent advancements in discovery.

Some references document the provenance of information that was used to inform the inception, design, or execution of the main experiments of the new study. These references are the ones that indicate some degree of causality. In other words, the prior work was used in such a way that the experiments and results would have been different (or not even performed) without the prior knowledge being referenced. In the context of biomedicine, this knowledge flow is thought to drive the discovery of fundamental principles of biology, the causes of disease, and effective therapies (1). These are the most important for researchers, funders, and administrators to identify in order to understand the past and future progression of biomedical research.

However, other references are discursive in nature. These help to place new results in the context of other studies in the field. They can showcase the novelty of the new discovery. In some cases, such references are perhaps aimed at persuading reviewers and editors to accept the paper for publication. These references are important to the business of science and the process of communicating, but lack the same causal nature of those that described prior work that was built upon. A small proportion of references track ideas that the authors consider, but then reject. Such ‘negative’ citations are estimated to be < 10% of the references in the literature (7–9), a figure that may vary in degree by field. It should be noted that these citations are still a marker of scientific influence, since they are thought to accelerate the self-correcting nature of science (7). Finally, some references are strategic in nature, designed to game citation metrics, or draw reader attention to the author’s prior work. Often, self-citations are viewed through this lens (10), although this interpretation is debatable (11, 12). The omission of these classes of references could, in principle, be an avenue for improving signal-to-noise in tracing the provenance of important ideas that contributed to human health through advancing clinical research.

Given the interest in identifying references that represent a causal relationship, and are well-aligned with the interpretation of a scientific article having impact (13–16), we asked whether it is possible to train a scalable model to identify the likelihood that a reference is causal in nature. Prior work has focused on identifying the type of discourse surrounding the reference in text (known as the “citation context”), as a proxy for author intent. Such models have made significant advancements in recent years (17–22).

However, approaches using citation contexts face two major limitations. First, it is unclear the extent to which different classes of discourse reveal a causal relationship between the citing and referenced work. Second, and most seriously, they require full text as an input and most articles are still not open access. This limitation means that even if hypothetical models were perfect, they could only capture a minority of causal citations at best. In biomedicine, the PubMed Central repository of public access articles captured 7 of the 32 million biomedical research articles in PubMed in 2021. This means that approximately 80% of causal citations in biomedical research would be overlooked even with a perfectly accurate model.

Here, we use information that is publicly and widely available to overcome this limitation of coverage. We asked whether information embedded in the combination of title/abstract text, article metadata, and the public domain citation graph (2, 23), contains enough information to indicate the likelihood that a reference is causal in nature. We leverage a data resource that has only recently become available at scale in biomedical research: biomedical preprints indexed in bioRxiv. When compared with the published version of the same manuscript, these reveal important information about the (non)causality of some references because these were added after the main body of experiments were conceived, designed, and executed, and therefore in general did not contribute to these formative steps of the research study. This interpretation is strongly supported by empirical work showing that authors deem the references they added during review to have been unimportant to these formative stages of research (24). This should not be surprising, since the main mechanisms by which authors might fail to include such papers in a pre-review reference list are most likely (1) authors were unaware of these papers, in which case it is hard to argue that direct knowledge transfer occurred at all, or (2) the authors were aware of these papers yet chose not to cite them. Here, we describe a model that is trained to identify such putative non-causal references, and also learns about the characteristics of causal references in the process. We show with classes of citations that are known a priori to be causal that the prediction scores of this model can be interpreted as a type of causal uncertainty. We then apply our model to references from clinical articles and studied the resultant causal uncertainty scores to gain insights into the efficacy of government funding (defined, for US biomedical research in this paper, as NIH funding for research articles indexed in PubMed) in stimulating the causal flow of information from basic to clinical research.

## Results

### Identification of non-causal citations

Large-scale identification of the subset of citations that are likely to be causal in nature is not a solved problem (25–28). However, researchers have identified subsets of citations that are *a priori* less likely to be causal in nature. Survey data asking researchers about the importance of the references that they themselves added to their publications shows that those added late in the writing process are far less likely to be important for the inception, design, or execution of the main body of experiments (24). Although it is possible for such citations to have contributed substantially to the small number of additional experiments requested during peer review, citations added after the writing and submission for publication of a largely complete scientific manuscript are therefore highly questionable as to their causal contribution to a scientific project.

Until recently, it was not possible to identify in the context of biomedicine which citations were added after the writing and submission of a scientific publication. However, biomedical researchers’ recent embrace of preprints has made this possible. The National Institutes of Health defines preprints as “a complete and public draft of a scientific document” and encourages their use and citation (29). There is evidence that biomedical researchers at present use preprints as largely complete scientific documents. The similarity of preprints and their later peer-reviewed version is high (30). Peer reviewed publications on COVID-19 rarely change the sentiment of their claims compared to their preprint version (31). Preprints describing clinical trials on COVID-19 did not change their conclusions during peer review (32). Finally, primary data from preprints are remarkably stable as they undergo peer review (33, 34).

Together, these findings support the theory and evidence that citations added in peer review, in general, are unlikely to have contributed to the inception, design, and execution of the core experiments constituting a study, which we refer to in this manuscript as the definition of a ‘causal citation’ (24). It should be noted that citations added in peer review may be related to additional experiments undertaken in response to reviewer criticism. However, the community is divided on the utility of these additional experiments relative to the time and expense of conducting them (35), and these results are often relegated to supplemental materials (36). Therefore, at a minimum, the *uncertainty* that these citations made a causal contribution to the main body of work (hereafter referred to as *causal uncertainty*) is very high.

To index citations that were present in biomedical preprints, we downloaded the full set of biomedical preprints from bioRxiv linked to peer-reviewed versions, identified using the Europe PubMed Central application programming interface. Citations were resolved with the Hydra citation resolution service (4). We then cross-referenced these preprint citations with those appearing in the peer-reviewed, published versions of the same paper using the NIH Open Citation Collection (2). In general, 11.6% of citations that were present in the peer reviewed version of the paper were added during review.

The causal uncertainty of references added during review may be high, but most publications do not have preprints with which to compare reference lists. However, it may be possible to identify patterns in the citation network and metadata of a paper and its references that represent citations with higher causal uncertainty. This problem is challenging because citation dynamics are noisy and therefore difficult to analyze. However, citation dynamics do follow certain mathematical principles (14, 15). By combining measures of article content with measurements of citation dynamics, it is possible to detect enough statistical regularity to predict, years in advance, outcomes like knowledge transfer from basic to clinical research (3). Therefore, it stands to reason that there may be enough statistical regularity in an article’s metadata to identify which citations have a high degree of causal uncertainty on that paper’s core experiments using machine learning.

Many machine learning models, in addition to learning patterns in the training data to identify the outcome measure that they were assigned, also learn enough about the underlying data structures to transfer that knowledge to answer other research questions. For that reason, we asked whether machine learning models developed to identify citations with high causal uncertainty could also identify citations that the scientific community expects *a priori* to be causal in nature, i.e. those with low causal uncertainty (more likely to represent a causal citation).

### Feature space

In order to generate a feature space in which a machine learning classifier could detect patterns in the content network that are associated with a citation being added during peer review vs. appearing in the original preprint, we considered information in three categories. The first category is metadata about the citing and referenced articles themselves. These included publication years of each paper, the degree of Human, Animal, and Molecular/Cellular Biology focus for each paper (3, 37), whether the citing and referenced paper appeared in the same journal, and whether the citing and referenced papers are primary research articles as defined in the NIH iCite tool (38).

The second category of information we included was information about the citation network and relationship between the citing and referenced articles. To encode this information, we generated measures that reflect size of the network, its growth rate, and particular types of citation relationship. First, we included information about the citing and referenced articles’ Relative Citation Ratio (RCR), which is a field- and time-normalized metric of an article’s citation rate (39, 40). Since its dissemination, RCR has been used by NIH in its portfolio analysis supporting science policymaking, strategic planning, and as evidence of good stewardship of taxpayer funds in U.S. Congressional Appropriations hearings supporting increases to the NIH budget (41–44). We also included information about ranking of these RCRs relative to the NIH-funded publication portfolio (here termed the RCR Percentile, but also referred to as Weighted RCR).

Previous work showed that highly cited papers are more likely to be the recipients of causal citations as identified by authors (24). We confirm the inverse, that papers with high causal uncertainty have lower ranked citation rates in general; the RCR percentile of papers where the reference was originally found in the preprint was 2.4% higher than for papers whose reference was added during review (p < 0.001, Wilcoxon Rank Sum test). This is not because older papers have had more time to accrue citations. The relative age of citations added in peer review was lower (6 year gap) than for citations originally found in the preprint version (7 year gap, p < 0.001, Wilcoxon Rank Sum test). In addition, RCR percentile time-normalizes its measure of scientific influence, so this comparison would not be affected by differences in publication age. Instead, references to papers added in peer review appear to have notably lower citation influence when adjusting for field and time.

We considered second-order information about the citation relationship as well (that is, two steps away from the target publication on a citation or co-citation graph) and included three additional measures of the citation network. First, we identified the number of other articles in the citing paper’s reference list that also cited the referenced paper (Figure 1a). This could be interpreted as a measure of the authoritativeness of the referenced paper, since these other related works deemed it important enough to cite. Next, we identified whether a paper published after the two papers in question has subsequently cited both the citing and referenced papers (i.e. indicating the citation in question is both a direct citation and a co-citation). A direct and co-citation relationship (Figure 1b) could be an indication that the two papers have been built upon and utilized together in a later scientific project. Third, we considered the local citation network of the full set of articles referenced by the citing paper, and asked whether the referenced paper falls in the largest connected component of that network (Figure 1c). Falling outside that largest component might convey information that the referenced paper is less related to the main body of related work.

**Figure 1.**
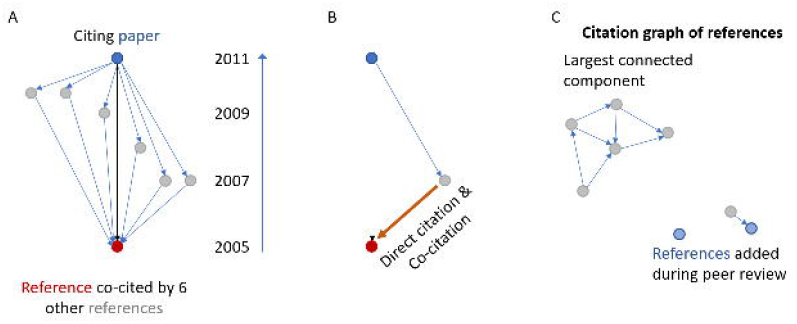
Local citaiton network features that may hold predictive power for citaiton causality. A) Illustration of references (grey) of a citing paper (blue) that all co-cited the target referenced paper (red) found in the same citing paper’s reference list. In this case, 6 other papers from this reference list (grey) co-cited the referenced article (red), so the count is 6. B) Illustration of a direct citation that is also a co-citation. The blue paper cites both the other citing (grey) and referenced (red) articles, making this orange direct citation link also a co-citation. C) The local citation network of the papers found in the reference list. Many referenced papers appear in the largest connected component, but in this illustration the two papers (blue) that were added to the reference list during peer review are not part of this component.

The third and final category of information we included was the content similarity of the citing and referenced articles. Recent advances in natural language processing models have advanced the comparison of the similarity of scientific documents. A recently developed deep learning model, SPECTER (22), has also integrated network-aware contextual information during training to teach the model whether the documents are also likely to have a direct citation linking them in addition to being semantically similar. This pre-trained deep learning model, when used for inference, jointly provides information about the semantic and citation network similarity. We generated features using SPECTER that encode cosine similarity information for the citing- and referenced article pairs, as well as information about the similarity of the citing article to the rest of its reference list (mean similarity and standard deviation), and the referenced article to the other papers in the reference list (mean and standard deviation). Finally, to capture information about the cohesiveness of the overall reference list, we generated features encoding the similarity of all of the references to one another (mean and standard deviation).

### Model training and validation

Collectively, these features comprehensively encode information about the citing and referenced articles, their relationships to one another, and the overall context of the other articles found in the reference list. These features have the advantage of being available at the time of publication and not reliant on additional information only disclosed after publication, as later forward- citations would. We assembled balanced training and testing datasets where each instance represented information about the content network regarding a single citing/referenced article pair. For the outcome measure, we applied a binary classification indicating whether this was a citation that was added in peer review (positive) or retained from the original preprint (negative). Some features that were initially promising were empirically tested and dropped from the final modelling process (see Supplemental Materials). We tested random forests, logistic regression models, support vector machines, and extreme gradient boosting (XGBoost). XGBoost (45) was selected for the remainder of the project because it had the highest predictive accuracy of these algorithms.

Overall, the model achieved an F1 accuracy score of 0.7 (Figure 2) on the test dataset, compared to a chance rate of 0.5. This corresponds to a reduction in uncertainty by approximately 40%. The distribution of prediction scores on the test dataset after training is balanced, but bimodal rather than uniform, weighted toward the ends (Figure 2c). This degree of accuracy is notably lower than models that take advantage of the sentence text in which a reference was cited (and the number of times a reference was cited in a manuscript). The main limitation that we seek to overcome is not a lack of predictive power given such a citation context. Instead, we seek to overcome the massive coverage gap: quantitative measures that rely on citation contexts by definition omit the vast majority of citation links because they are not open access. In this overlooked space that constitutes the vast majority of the references of the scientific literature, there is an opportunity to shed light into which citations are among the most- or least-likely to represent causal information transfer.

**Figure 2.**
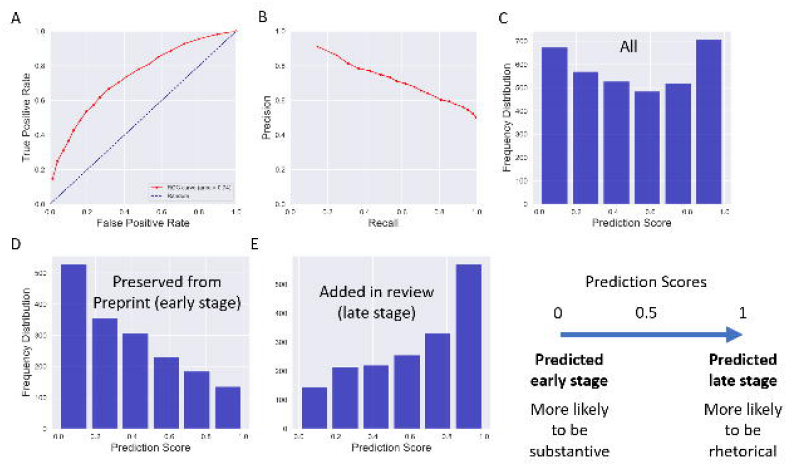
Model accuracy. A, B) Precision/recall graph and ROC curve of prediction scores on out-of-sample test data. C) Distribution of prediction scores on a balanced set of out-of-sample positives and negatives. D) Distribution of prediction scores for out-of-sample positives. E) Distribution of prediction scores for out-of-sample negatives.

Feature importance scores are shown in Table 1. Out-of-sample positives received a higher prediction score of 0.63 on average, while out-of-sample negatives received an average prediction score of 0.37 (p < 0.01, Wilcoxon rank sum test, and Figure 2d-e). This is important because it is possible for a prediction system to have predictive power mainly for identifying positive values, while performing closer to chance at identifying negatives. If that were the case, it would indicate that the model had learned some predictive properties of papers added in peer review, but little else. The low distribution of prediction scores for the out-of-sample negative samples indicates that the model is learning patterns about citations that are possibly causal as well.

**Table 1.** Feature importance scores from the trained model. See Methods for full feature descriptions. Gain, fractional contribution of features, indicating a more predictive feature. Cover, relative number of observations related to the feature. Frequency, a count of the relative number of usages of a feature within trees.

Beyond accuracy testing, we asked whether there were other tests of model validity that could confirm or disconfirm that the model has learned how to predict citations with high causal uncertainty. We first identified a class that is enriched in citations with high causal uncertainty. First, we turned to citations that were present in the preprint version of a paper but were excluded in the published version. One caution about this dataset is that there is more than one reason that the citation could be dropped. First, an author may have deemed the citation unimportant. In this case, the interpretation that it was unlikely to be causal in nature is relatively straightforward.

However, a second reason the citation might not appear in the published version has to do with an idiosyncrasy of the data processing pipelines of preprints vs. publishers. bioRxiv, the source of our preprint citation data, includes supplemental references in its online structured citation data. Many publishers do not, and so the citations may not have been dropped at all, but rather were in the supplement, where citations are not indexed for the publication version at many journals. Nevertheless, because both mechanisms can lead to the apparent removal of the citation, there should be more non-causal citations in these citations than in a random sample of citations. Thus, these citations should have higher than average prediction scores if the model is learning features correlated with high causal uncertainty. Figure 3a shows that this is indeed the case (p = 0.0011, Wilcoxon rank sum test), supporting the hypothesis that the model identifies such correlates.

**Figure 3.**
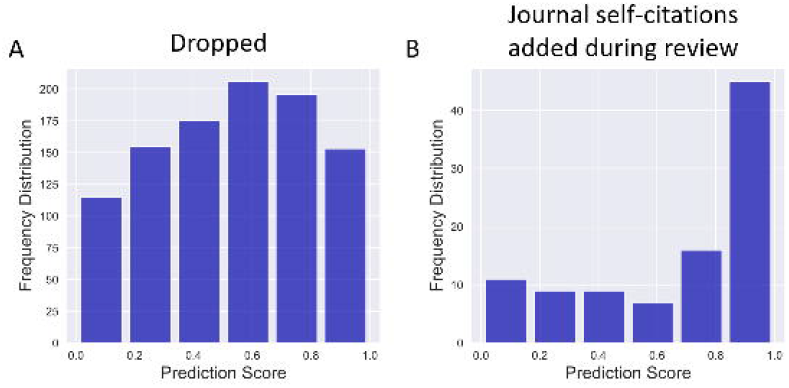
Prediction scores are higher for citations that *a priori* carry higher causal uncertainty. A) Distribution of prediction scores for preprint references that were dropped in peer review and no longer appeared in the published version. B) Distribution of prediction scores for possibly coercive citations, those that were added in peer review and cite the same journal that the paper was submitted to.

We next turned to a class of citations that has gathered much scrutiny because of their potential for gaming of citation statistics. These are citations that were added in peer review and cite earlier papers from the same journal. These citations are highly suspect; editors insisting on the addition of such citations, in order to game journal-level citation metrics like the Journal Impact Factor, before acceptance have been termed “coercive citations” (46). Note that authors need not wait for editorial feedback before adding such non-causal citations. They may instead anticipate more favorable editorial decision-making if the citations are added, and preemptively add these (46). Thus, if the model has learned about causal uncertainty, these should also receive high scores. We examined out-of-sample citations to the same journal that were added in peer review. Notably, these putative coercive citations received the highest prediction scores for being non-causal of any class of citation we examined (p < 0.001, Wilcoxon rank sum test and Figure 3b), even the out-of-sample positives.

### External validation

Since these results show that the trained model has learned to predict the citations that we have *a priori* reason to believe are less likely to be causal in nature, we turned our attention to the question of whether it learned about citations that are more likely to be causal as well. We identified five classes of such citations as tests of the trained model’s external validity.

First, we relied on the highly structured regulatory framework surrounding drug development to identify causal citations. In the United States, this framework constrains the generation of clinical knowledge and applies order to this process. Clinical trials proceed in phases that test first for safety and then for efficacy, progressing to larger trials that build off the preceding trials. Therefore, to comply with clinical regulation, later-stage clinical trials for a given drug must by definition draw on previous, lower-stage clinical trials assessing that same drug for treating the same disease. These citations from later-stage clinical trials to earlier stage clinical trials for the same drug therefore are among the most likely citations to represent causal knowledge transfer. Our model was trained to identify citations with high causal uncertainty (i.e. those least likely to have contributed to the inception, design, or execution of the main body of a paper’s experiments). If the model has learned that putatively causal citations are, in many respects, the opposite, the prediction scores for this opposite class of citations should have low values. Figure 4a shows that this is the case (p < 0.001, Wilcoxon rank sum test). Thus, the prediction scores can in some sense be interpreted as an estimate of causal uncertainty.

**Figure 4.**
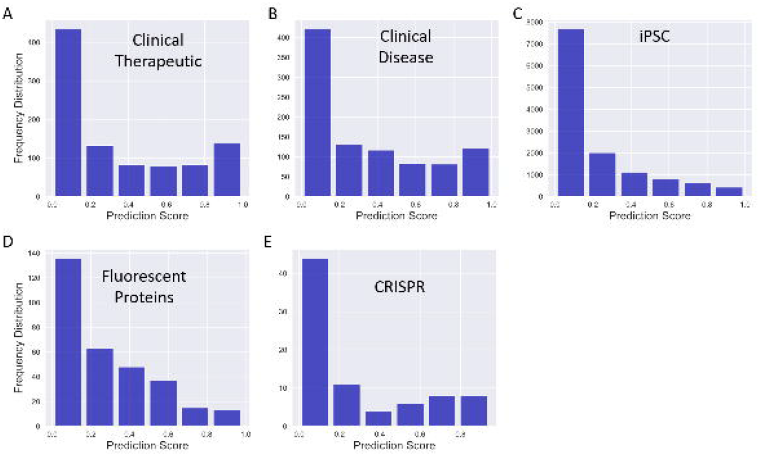
Prediction scores are lower for citations that *a priori* are known to be causal, and can be interpreted as a measure of causal uncertainty. A) Prediction scores for citations from higher- phase clinical trials to preceding, lower-stage clinical trials matched by the drug therapeutic. B) Prediction scores for citations from higher-phase clinical trials to preceding, lower-stage clinical trials matched by the disease being treated. C) Prediction scores for citations from iPSC papers to the paper describing the seminal iPSC methodology. D) Prediction scores for citations from papers using fluorescent proteins to the paper describing novel fluorescent proteins for use by the research community. E) Prediction scores for citations from CRISPR papers to the paper describing the seminal CRISPR gene editing technology. F) Visual interpretation of model prediction scores. Higher scores (i.e. those most similar to references added during peer review and therefore least likely to have informed the inception, design, or execution of the main body of experiments) are less likely to represent causal citations, while those with the lowest scores are validated as being more likely to represent causal citations.

We next asked whether the model’s external validity extended to clinical trials studying the same disease rather than the same drug. While perhaps less tightly ordered around the regulatory framework governing drug development, we reasoned that this class of citations, when they occur, are likely to be enriched in causal citations because of their shared goal of treating a common disease. This is especially true because some clinical trials are structured to compare novel treatments with the established standard of care, where the regulatory framework encourages knowledge flow within the same disease topic but across different drugs. Here, too, the model gave low prediction scores (Figure 4b), consistent with a successful prediction (p < 0.001, Wilcoxon rank sum test).

We next turned our attention to fundamental research with a study that rapidly emerged into its own research topic. Rather than drawing upon the US regulatory framework, we next drew upon acknowledged contributions to the scientific literature by Nobel prizewinning research. In the mid-2000’s, human embryonic stem cell (hESC) research was a novel and promising approach to modeling human biology, contributing to therapeutic development, and being used as a possible therapeutic agent themselves. At that time, the number of hESC papers in PubMed was growing exponentially. However, there was unmet demand from the scientific community and the public for sources of pluripotent stem cells that carried fewer ethical concerns about abortion and might be subject to less administrative oversight and restriction.

In 2006, the Yamanaka lab produced induced pluripotent stem cells (iPSCs) from adult, rather than embryonic, cells (47). This discovery was followed shortly thereafter in 2007 by production of human iPSCs by the Yamanaka and Thompson labs (48). iPSC research rapidly spread through the cell biology community, and by the mid-2010’s eclipsed hESC research (which had started to decline) and is still growing rapidly. Citations from the later iPSC papers that used or improved upon the original study’s methodology can be inferred to be causal in nature. We identified iPSC papers that cited the original iPSC article (47), and asked whether these, too, received low prediction scores. Figure 4c shows that this is the case (p < 0.001, Wilcoxon rank sum test). Thus, this model can identify references, with low prediction scores, that are known *a priori* to have contributed to the genesis of a new research topic.

Studies that spawn a research topic are rare, and we asked whether enabling technologies that spread rapidly into other established topics are also identified. The development of fluorescent proteins fits this description, although earlier works describing the Green Fluorescent Protein (GFP) might be considered to have emerged into its own scientific topic. This technology enabled the tagging of individual proteins within the cell, so that their subcellular distribution could be quantified with a genetic, rather than chemical, approach. For that reason, we examined later-stage fluorescent protein development, when fluorescent protein research had already emerged as a topic of scientific inquiry, but new fluorescent proteins with unique spectral properties represented an enabling technology for other, already established fields of science (49).

In order to identify suitable citations, we turned to the methods section of articles with free full-text in PubMed Central. This is because, unlike the iPSC topic, we sought papers that were not necessarily studying fluorescent proteins per se, but rather were more likely to use them as a means to an end. Open science databases such as OpenCitations and COLIL have parsed the citation context (50, 51), which is the sentence in which a reference is cited, alongside the section of the article in which the citation is found. We queried references appearing in the Methods section of articles in citation contexts containing names of specific fluorescent proteins that were developed after the emergence of fluorescent proteins as a research topic, but that were still widely used and disseminated in other fields. Like the other classes of citation used for external validation, these are expected to receive low prediction scores if the model is successful. Figure 4d shows that these fluorescent protein citations also receive low prediction scores (p < 0.001, Wilcoxon rank sum test). Citations to enabling technologies in two contexts, then, can be detected, without respect to whether the cited article launched a new topic of scientific inquiry.

Finally, we asked whether citations to innovations with particularly high commercial potential are also identified. This is not to say that the commercial potential of iPSCs and fluorescent proteins were not high, but some technologies are known *a priori* to have especially high commercial potential. Broadly effective genome editing technology is one of these. In 2013, Doudna and Charpentier described a novel gene editing approach using the CRISPR/Cas9 system (52). Subsequently, patent rights for its application were sharply disputed (53). To test whether subsequent citations from papers employing this technique are identified by our model, we again extracted citation contexts from citing papers and included citations to the CRISPR/Cas9 article that appeared in the Methods section. This final class of citations that are thought to be causal in nature also received low scores by the model (p < 0.001, Wilcoxon rank sum test and Figure 4e). Thus, the trained machine learning model seems to have identified not only that citations with high prediction scores are less likely to be causal citations that contributed to the inception, design, or execution of the main experiments in a study (e.g. similar to those added during review), but also gives the opposite prediction scores (i.e. low absolute prediction values) for known causal citations (Figure 4f).

### US Federal support of fundamental knowledge cited by clinical studies

One pressing question in the biomedical research literature is how to more effectively facilitate the generation of knowledge that will drive later clinical discovery. However, before applying our causal uncertainty scores to address this question, it is first necessary to understand the overall dynamics of bench-to-bedside translation in recent years. We focused on the distinction between federally funded vs. non-federally funded research within the United States because science funding is a major policy lever easily accessible to lawmakers for advancing science, and the value of government funded science is under constant scrutiny. Using the Human, Animal, and Molecular/Cellular Biology classification initially developed by Weber (37) can help to identify the domain of science for each paper. Human publications, which are conceptually the closest to clinical research, while Animal and especially Molecular and Cellular Biology research are conceptually farther. These can be visualized on the Triangle of Biomedicine (3, 37), which places articles into a translational space that visually depicts where articles are found in this space. Human research articles characterize the majority of biomedical studies. By contrast, NIH-funded research has a more basic research focus (Figure 5a), although this difference is closing over time (Figure 5).

**Figure 5.**
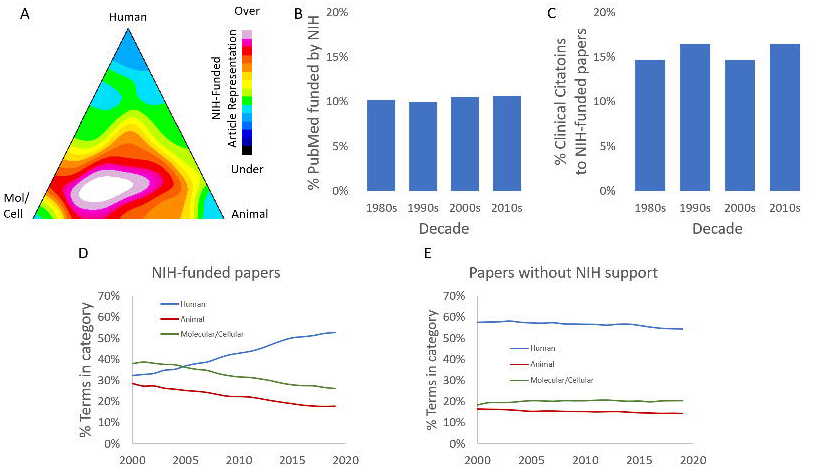
US Federal support for Human, Animal, Molecular/Cellular Biology research. (A) Density graph of over- and under-representation of NIH-funded articles in different domains of the Triangle of Biomedicine. NIH-supported papers are overrepresented in Molecular/Cellular Biology and Animal research, and comparatively underrepresented in Human research. (B) NIH-funded articles comprise ∼10% of the biomedical literature. (C) NIH-funded articles typically receive 15-16% of citations from clinical research articles. (D) Shifts of the domains of research funded by NIH over time. (E) The non-NIH-funded literature has a strong and stable human focus.

It stands to reason that the more basic-research favored for federal funding might be less-well cited by clinical articles, due to the larger conceptual distance to applied clinical research. Extending this line of reasoning, it may be that those clinical citations that exist might be less likely to represent causal knowledge transfer because of the large conceptual gap between basic and applied clinical research. Over the period we studied, NIH-funded scientific papers represented 10.6% of biomedical publications (Figure 5b). We therefore asked whether US federally funded research articles represent more or fewer than ∼10% of clinical citations as a first test of the representation of US federally funded biomedical research in clinical citations. NIH identifies clinical citations as citations from clinical observational studies, trials, or guidelines in its iCite web service (3). NIH-funded articles comprise 15.7% of papers cited by clinical articles (Figure 5). Despite the more basic research focus, NIH-funded articles receive nearly 50% more clinical citations than might be expected based on their fraction of the literature. However, because of the limitations of citation analysis, this result does not necessarily reveal the over- or under-representation of *causal* knowledge flow from federally- supported to clinical articles.

### Causal uncertainty in government-supported bench-to-bedside translation

To address questions about *causal* clinical citations, we first had to operationalize this concept for the purposes of this study. In our external validation studies involving citations that we have reason *a priori* to believe represent causal knowledge flow, like iPSC and progressive phases of clinical trials involving the same drug, these generally scored below 0.5 on predictions from our model. Likewise, clinical citations to NIH-funded articles had a similar distribution of prediction scores (Figure 6a). We asked if this pattern changed when examining clinical citations to basic NIH research, given the greater conceptual distance between basic and clinical research. We observed that these clinical citations also had a low distribution of prediction scores (p < 0.001, Wilcoxon rank sum test Figure 6b), and also resembled the external validation datasets that are *a priori* likely causal citations.

**Figure 6.**
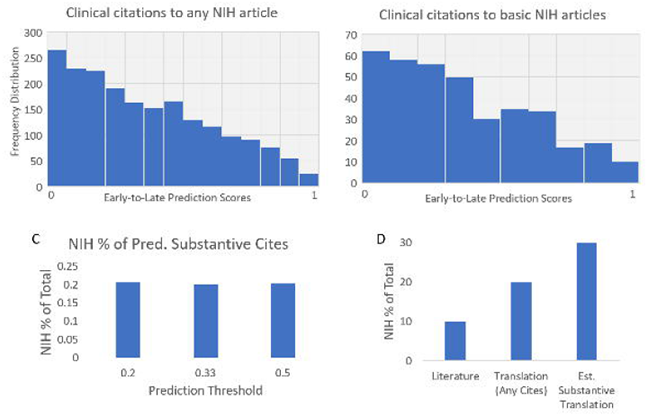
Government support overrepresented in estimated causal knowledge flow from basic to clinical research. (A) Distribution of causal uncertainty scores for clinical citations to any NIH-funded articles. (B) Distribution of causal uncertainty scores for clinical citations to basic research papers funded by NIH (referenced papers have Human scores below 0.5). (C) NIH- funded papers receive 20% of citations with causal uncertainty scores below threshold, regardless of the threshold chosen. (D) NIH-funded papers are overrepresented in clinical-to- basic citations (20%, no filtering on causal uncertainty, “Translation (Any Cites)”). NIH-funded papers comprise nearly a third of basic research articles that receive clinical citations with a causal uncertainty score below 0.2 (“Est. Causal Translation”).

Based on the response characteristics of the model’s prediction scores and the external validation studies shown in Figure 4, we reasoned that citations with low prediction scores are likely to be enriched in causal citations. To be clear, we do not assert that each individual citation with a low causal uncertainty score is a causal citation. Instead, given the F1-score of 0.7, we estimate that a comparable proportion of causal citations are likely to be found in the group of citations with a low causal prediction score. We therefore refer to this group of citations as *estimated causal citations*. Using this initial threshold as an operational definition of estimated causal citations from clinical articles, we found that NIH-funded publications comprised 20% of such citations, twice as high as the representation of NIH-funded publications in the scientific literature (Figure 6). Limiting to the estimated causal clinical citations thus reveals that, compared to the 15% of clinical citations received by NIH papers overall (Figure 5), US federally funded studies are overrepresented among those citations most likely to have driven causal knowledge transfer. Furthermore, this estimate was invariant to the particular threshold used. NIH-funded publications received 20% of the total likely causal clinical citations using more stringent thresholds of 0.33 and 0.2 as well (Figure 6c), the latter of which was the most stringent and the most concentrated in causal citations from our external validation (see Figure 4), and therefore used in the next analysis. Thus, independent of the particular threshold used, federally funded biomedical research is overrepresented twofold among clinical citations estimated as likely to be causal by our model.

We next asked whether this finding extended to clinical citations to basic research articles rather than NIH-funded articles overall. This is because, although less Human-oriented than the literature as a whole, the NIH portfolio is increasingly focused on Human-focused research articles (Figure 5). Previous work has identified research that is less than 50% human focus as enriched in basic research articles (3, 37). Using that definition, we found that NIH-funded research articles comprised 30% of estimated causal citations from clinical to basic research articles (Figure 6). Thus, although NIH-funded work comprises only 10% of the literature, it represents an estimated 30% of the basic research literature that represents likely causal knowledge transfer into clinical research (p < 0.001, Chi-squared test for probabilities).

## Discussion

An important applied goal of biomedical research is to generate knowledge that can advance human health. US Federal science funding is among the most prominent policy levers for stimulating basic research knowledge that could inform new therapeutic development. Citations represent knowledge flow from the referenced article to the citing one, and can be used to trace the movement of basic research ideas into such downstream applied research. Recently we developed a large-scale database of citations from the clinical literature to the rest of biomedical science, which for the first time can be used to address these kinds of questions (2, 3, 38). However, the presence of a citation does not reveal how that knowledge was put to use. Some references are exclusively discursive in nature, and may not have been meaningfully built upon in the citing study. This is a known limitation of citation analysis, which makes the tracing of knowledge flow in bench-to-bedside translation noisier than would otherwise be the case. While citation serves as an acknowledgment of intellectual contribution to the specific research, not all citations are intended to perform the same task. For example, some citations are intended to provide the context of the problem or discuss seminal works in the field and such citations are mostly found in the introduction. Citations in the related work mostly discuss previous works on the same topic and the limitations of those works. Citations in the experiment sections are mostly related to baseline comparison and contribution analysis. These kinds of citations are important to provide the readers with enough context and knowledge on the topic and bring them to the same page as the authors. However, these citations do not necessarily have intellectual impact specifically with respect to the inception, design, or execution of a citing study. Many of those might be discovered at a much later stage, therefore, it will be misleading to claim those as the causal citation in this narrow context.

Our results demonstrate that it is possible to use information about local network structure, deep learning semantic representations of text similarity, and article metadata to discern a signal of citation causality from the noisiness of citation dynamics. We took advantage of knowledge from preprints and their published versions to identify a class of citation that is, a priori, highly unlikely to have been used to inform the inception, design, or execution of the main experiments of the new study: those added during peer review, after the main body of work has been completed (24). We then trained a model to predict when the certainty is *a priori* low that a referenced paper contributed to these early stages of research development and execution. We found that citations that are known historically to have been causal in nature receive low causal uncertainty scores. Finally, we apply this model and find that federally funded publications are vastly overrepresented in the population of basic-to-clinical citations with low causal uncertainty scores. This population of citations is likeliest to represent causal knowledge transfer in a bench-to-bedside translation context, indicating that federal support for biomedical research is a highly effective tool for stimulating this process.

This study has important limitations. First, because of the scope of this work and the data utilized (2, 23, 38), it is limited to biomedical research fields. It is possible that the model efficacy at identifying causal citations would change if expanded to other fields. Second, since preprint data are relatively recent in biomedicine, it may be the case that this modelling approach is most effective at making predictions about recent citations, rather than historical ones from decades past. This is especially salient because the later publications analyzed contain COVID-19 publications. It is not known if citation or semantic relationships shifted during the pandemic. Further, it is not known whether the network structures and semantic representations used for feature generation drift over the scope of decades. However, it is our anticipation that further research into identifying causal citations without full text will increase the accuracy and coverage of information about causal knowledge transfer and identify effective methods to combine the approaches reported here with the rich information for citations where contexts are fully available.

Evidence from the literature supports the interpretation that citations added during review are unlikely to have informed the inception, design, or execution of the main experiments. This is because articles do not change much during the review process, leaving little scope for this stage of science communication to effect change. With an intention to rapidly and openly share research findings with the community, biomedical researchers have increasingly accepted preprint servers. Several studies across different research domains compared preprints with their peer-reviewed counterparts to analyze the credibility of preprints as well as the effectiveness of peer-review. One study (54) manually compared the quality of reporting of both independent and paired preprint and peer-reviewed samples from bioRxiv and PubMed. Their findings suggest the quality of reporting in preprints and peer-reviewed articles is within a comparable range although the peer-reviewed articles have a slightly better quality of reporting on average. Dirnagl et al. analyzed the preprints recording the early phase of the COVID-19 pandemic and their subsequently peer-reviewed versions (55). They compared the number of figures, panels, and tables and found no difference on average, suggesting that very few new experiments or analysis were conducted during the peer-review. Moreover, the abstracts and conclusions of the preprints experienced very minor changes during review. However, Xu et al. (56) found that preprints that were published performed better in altmetrics than those preprints that were never published, indicating important differences in usage among the scientific community between preprint-only manuscripts vs. those published. Other studies analyzed the text-similarity and semantic features of preprints and their peer-reviewed counterparts and the findings are consistent across different fields (57, 58). The semantic analysis of the title, abstract, and body of the preprints in arXiv and bioRxiv found little notable difference with their peer-reviewed versions (57). Comparison of preprints and their peer-reviewed versions on COVID-related research also found coherence in the reported results (54, 57, 59), extending the findings of these studies. In addition, analysis of the primary data constituting the evidence base suggests that estimate values in published papers are nearly identical to those in preprint versions (32, 33). In addition, there are not measurable differences in peer review evaluations of article quality between preprints that are published vs. those that remain unpublished (33). Studies that examined non-COVID preprint-publication pairs drew similar conclusions (60–64). These studies support the use of preprints, as one important component of the scientific literature, in downstream research development and decision-making.

The methods explored here should not be viewed as an alternative for citation context-based approaches for identifying important citations (17–22). These two approaches, although aligned in their goals, utilize non-overlapping information, and the insights from each type of model can be combined to further reduce uncertainty about citation causality. Instead, the approaches used here can be used to address the large coverage gap in open access to free full text, and dramatically expand the scope of citations that can be examined from the context of causality and importance. We show that it is possible to generate predictions about citation causality even without access to full text citation context.

Finally, our results indicate that federal support of science funding is a powerful lever to translate basic knowledge into clinical discovery. These results extend earlier work suggesting that most new therapeutics, typically privately funded, build on earlier federally funded work that formed the basis for new drug development (65). Our results indicate that this prior federally funded basic research is among the most likely to have fueled causal knowledge transfer to later clinical research.

## Materials and Methods

### Raw data & code availability

Preprints that have been published were identified through the Europe PMC application programming interface, which matches the Digital Object Identifier (DOI) of the preprint and published versions of a paper. DOIs corresponding to bioRxiv preprints were identified and their full text, which includes a structured reference list, were downloaded in October 2020 (33). It should be noted that it is possible that more recent preprint-publication linkage may not yet have been indexed in Europe PMC due to possible reporting or data processing delays. For this reason, more recent preprint-publication links may be underrepresented in our sample. References were matched with the National Library of Medicine’s Hydra citation resolution service (4). We used the NIH Open Citation Collection to identify references from the final, published version of a manuscript (2, 23). We used publication-grant linkages from NIH ExPORTER to identify federally supported biomedical research articles (66).

Many of the derived features described below used data from the iCite database (e.g. Human, Animal, Molecular Biology scores (3), or the Relative Citation Ratio (39, 40) in its linear or percentiled form) (2, 3, 23, 40, 67). These are available at the Figshare data repository (67). Raw text for similarity comparisons are available at PubMed (68). Source code for the SPECTER and XGBoost libraries used in this study are available online (22, 69, 70).

### Feature descriptions

- `specter_cosine_sim`: SPECTER cosine similarity using the title and abstract text of the citing paper and that of the referenced paper
- `pub_ref_reflist_sd`: Standard deviation of the pairwise SPECTER cosine similarities using the title and abstract text of the referenced paper and that of all other papers in the citing article’s reference list
- `pub_ref_reflist_mean`: Mean of the pairwise SPECTER cosine similarities using the title and abstract text of the referenced paper and that of all other papers in the citing article’s reference list
- `pmid_reflist_sd`: Standard deviation of the pairwise SPECTER cosine similarities using the title and abstract text of the citing paper and that of all other papers in the citing article’s reference list
- `pmid_reflist_mean`: Mean of the pairwise SPECTER cosine similarities using the title and abstract text of the referenced paper and that of all other papers in the citing article’s reference list
- `reflist_reflist_sd`: Standard deviation of the pairwise SPECTER cosine similarities using the title and abstract text of all papers in the citing article’s reference list to one another
- `reflist_reflist_mean`: Mean of the pairwise SPECTER cosine similarities using the title and abstract text of all papers in the citing article’s reference list to one another
- `ref_year`: Publication year of the referenced paper
- `ref_rcr`: Relative Citation Ratio of the referenced paper
- `pub_rcr`: Relative Citation Ratio of the citing paper
- `cocited_by_ref`: Count of the number of other papers in the citing paper’s reference list that also cited the referenced paper
- `pub_pctile`: Percentile of the citing paper’s Relative Citation Ratio
- `ref_mc`: Molecular/Cellular Biology score of the referenced paper. This is the average of relevant Medical Subject Heading terms attached to this paper that fall into the Molecular/Cellular category
- `pub_mc`: Molecular/Cellular Biology score of the citing paper. This is the average of relevant Medical Subject Heading terms attached to this paper that fall into the Molecular/Cellular category
- `ref_a`: Animal score of the referenced paper. This is the average of relevant Medical Subject Heading terms attached to this paper that fall into the Animal category
- `pub_a`: Animal score of the citing paper. This is the average of relevant Medical Subject Heading terms attached to this paper that fall into the Animal category
- `pub_year`: Publication year of the citing paper
- `ref_pctile`: Percentile of the referenced paper’s Relative Citation Ratio
- `ref_h`: Human score of the referenced paper. This is the average of relevant Medical Subject Heading terms attached to this paper that fall into the Human category
- `pub_h`: Human score of the citing paper. This is the average of relevant Medical Subject Heading terms attached to this paper that fall into the Human category
- `in_lcc`: Binary flag for whether the referenced paper falls into the largest connected component of the local citation network of the papers in the citing article’s reference list
- `direct_and_cocitation`: Binary flag for whether the direct citation between the citing and referenced paper is also a co-citations; in other words, has another article been published after the citing paper that referenced both the citing and cited paper?
- `ref_is_research`: Binary flag for whether the referenced paper is a primary research article
- `same_journal`: Binary flag for whether the citing and cited papers are both published in the same journal
- `pub_is_research`: Binary flag for whether the referenced paper is a primary research article

### Machine learning & accuracy testing

Our outcome measure for prediction was a binary flag indicating whether a given reference had been added in peer review (True) or was found in the original preprint and carried over to the published version (False). Training data contained 440,000 instances of balanced positive and negative data, and accuracy statistics were calculated from a smaller holdout test set. XGBoost was used for the final model (70), although we also tested random forests, support vector machines, and logistic regression (each of which had poorer performance). The model achieved an F1 score of 0.7. Training and testing using the complete balanced dataset with 10-fold cross-validation yielded similar results. Our final test set comprised of 3482 citation linkages that were out-of-sample for training the model.

References that were found in the preprint version but not in the published version were omitted because of a difference in how such references are indexed. bioRxiv and medRxiv host both the primary reference list and references found only in the supplemental material, while many publishers do not deposit references from supplemental material into structured citation indices. It cannot be easily distinguished whether a reference is found in the preprint but not the published version due to being genuinely dropped, or whether it was part of the supplemental data and not covered by publishers. We therefore omitted such references from training data.

### External validation & analysis

To identify earlier stage clinical trials cited by later stage clinical trials for the same drug, we used PubChem to match drugs and PubMed Publication Type terms to identify which Phase (I-IV each citing/referenced paper was (68). To identify those trials studying the same disease, we matched based on the Disease Medical Subject Heading terms available in PubMed. To test the role of federal support on estimated causal translation on clinical citations, we randomly sampled 10,000 citations from clinical papers published in 2015-2019 for subsequent subsetting and analysis. Sample sizes for other external validation conditions were: Clinical progression of the same drug (955); clinical progression of the same disease (958), iPSC (12693), XFP (312), and CRISPR (81).

Some external validations required examining the location in the citing paper of the citation. Such citation contexts are available from OpenCitations and COLIL (50, 51); we used context data from the latter service. Only citations in sections that contained the case-insensitive string “methods” in the header were used for matching citations in the Methods section (e.g. “Methods”, “Materials and Methods”, or “Methods and Results”). Additional citations to the same paper in other sections of a paper did not exclude a reference from consideration as long as it was also cited in the Methods section.

## Acknowledgements and Funding Sources

BIH is funded through the Office of the Vice Chancellor for Research and Graduate Education at the University of Wisconsin-Madison and through funding from the Wisconsin Alumni Research Foundation. Disclaimer: this work was performed prior to TH joining the Centers for Disease Control and Prevention and should not be considered a research product of that agency. Disclosure: BIH has previously been supported as an NIH employee, as a trainee in the NIGMS Postdoctoral Research Associate Program, and from NIH grant F31GM080164 from NIGMS.

