## Supplementary material for "Predicting causal citations without full text": Table 1

| Feature | Gain | Cover | Frequency |
| --- | --- | --- | --- |
| specter_cosine_sim | 0.099 | 0.137 | 0.116 |
| pub_ref_reflist_sd | 0.098 | 0.149 | 0.120 |
| pub_ref_reflist_mean | 0.094 | 0.152 | 0.115 |
| pmid_reflist_sd | 0.081 | 0.096 | 0.071 |
| pmid_reflist_mean | 0.081 | 0.098 | 0.069 |
| reflist_reflist_sd | 0.077 | 0.101 | 0.069 |
| reflist_reflist_mean | 0.076 | 0.095 | 0.066 |
| ref_year | 0.067 | 0.019 | 0.050 |
| ref_rcr | 0.062 | 0.047 | 0.074 |
| pub_rcr | 0.035 | 0.023 | 0.032 |
| cocited_by_ref | 0.025 | 0.008 | 0.020 |
| pub_pctile | 0.025 | 0.018 | 0.021 |
| ref_mc | 0.024 | 0.005 | 0.028 |
| pub_mc | 0.023 | 0.005 | 0.021 |
| ref_a | 0.021 | 0.004 | 0.024 |
| pub_a | 0.019 | 0.004 | 0.017 |
| pub_year | 0.018 | 0.003 | 0.011 |
| ref_pctile | 0.018 | 0.020 | 0.020 |
| ref_h | 0.018 | 0.005 | 0.021 |
| pub_h | 0.017 | 0.004 | 0.016 |
| in_lcc | 0.010 | 0.002 | 0.006 |
| direct_and_cocitation | 0.006 | 0.002 | 0.006 |
| ref_is_research | 0.005 | 0.001 | 0.005 |
| same_journal | 0.002 | 0.001 | 0.002 |
| pub_is_research | 0.001 | 0.001 | 0.000 |
