## Supplemental Materials for "Predicting causal citations without full text"

### Supplemental Results

#### Feature space analysis

We first began with features extracted from three databases: PubMed, iCite, and Semantic Scholar. This initial model used the features described in the Methods: Feature Descriptions section (which derived data from iCite and PubMed), as well as four additional features derived from Semantic Scholar (Supplemental Table 1). This model, in addition to being more complex, also requires ingesting a third database, so its upstream data processing is also more complex. The four features derived from Semantic Scholar data each received feature importance scores near the bottom the importance ranking (Supplemental Table 2), so we explored creating a model without these (background, methods, results, authors_pmid_ref_overlap). Both models received similar F1 scores: 0.7 for the model including Semantic Scholar data, and 0.7 for the model using only PubMed and iCite data (see Results). A heatmap of the feature correlation matrix is shown in Supplemental Figure 1. Because these two scores were similar, we used the model without Semantic Scholar data to simplify the model and data processing pipeline.

| Additional feature | Description |
| --- | --- |
| background | Semantic Scholar’s estimate, based on their deep learning model of citation contexts, that this citation is a ‘background’ style citation |
| methodology | Semantic Scholar’s estimate, based on their deep learning model of citation contexts, that this citation is a ‘methodology’ style citation |
| result | Semantic Scholar’s estimate, based on their deep learning model of citation contexts, that this citation is a ‘result’ style citation |
| authors_pmid_ref_overlap | Measure of overlap between the authors’ published papers and those referenced in this paper (a measure of self-citation). Uses Semantic Scholar author publication profiles |

**Supplemental Table 1**. Additional features removed from final model.

| Feature | Gain | Cover | Frequency |
| --- | --- | --- | --- |
| specter_cosine_sim | 0.094354 | 0.132561 | 0.112403 |
| pub_ref_reflist_sd | 0.094085 | 0.148231 | 0.117254 |
| pub_ref_reflist_mean | 0.090423 | 0.148164 | 0.111646 |
| pmid_reflist_sd | 0.079526 | 0.098855 | 0.071388 |
| pmid_reflist_mean | 0.077426 | 0.09796 | 0.06851 |
| reflist_reflist_sd | 0.076695 | 0.098471 | 0.068842 |
| reflist_reflist_mean | 0.076608 | 0.097763 | 0.066528 |
| ref_year | 0.065068 | 0.019641 | 0.048819 |
| ref_rcr | 0.060807 | 0.046601 | 0.072988 |
| pub_rcr | 0.034135 | 0.022887 | 0.030774 |
| pub_pctile | 0.025242 | 0.017468 | 0.021312 |
| cocited_by_ref | 0.024214 | 0.007907 | 0.019668 |
| ref_mc | 0.024012 | 0.004573 | 0.028233 |
| pub_mc | 0.022668 | 0.004451 | 0.020438 |
| ref_a | 0.020082 | 0.003833 | 0.023841 |
| ref_pctile | 0.018071 | 0.018312 | 0.02027 |
| pub_year | 0.017811 | 0.003961 | 0.010814 |
| pub_a | 0.017683 | 0.003971 | 0.016499 |
| pub_h | 0.016961 | 0.003959 | 0.015513 |
| ref_h | 0.016905 | 0.004209 | 0.020117 |
| in_lcc | 0.00921 | 0.001785 | 0.00549 |
| background | 0.009112 | 0.002059 | 0.006828 |
| authors_pmid_ref_overlap | 0.008146 | 0.005574 | 0.005312 |
| direct_and_cocitation | 0.005883 | 0.001672 | 0.005696 |
| methodology | 0.005352 | 0.001125 | 0.003284 |
| ref_is_research | 0.005143 | 0.001319 | 0.004551 |
| same_journal | 0.001666 | 0.000801 | 0.001553 |
| result | 0.001614 | 0.001074 | 0.00093 |
| pub_is_research | 0.001097 | 0.000816 | 0.000499 |

**Supplemental Table 2.** Feature importance scores of the more complex model.

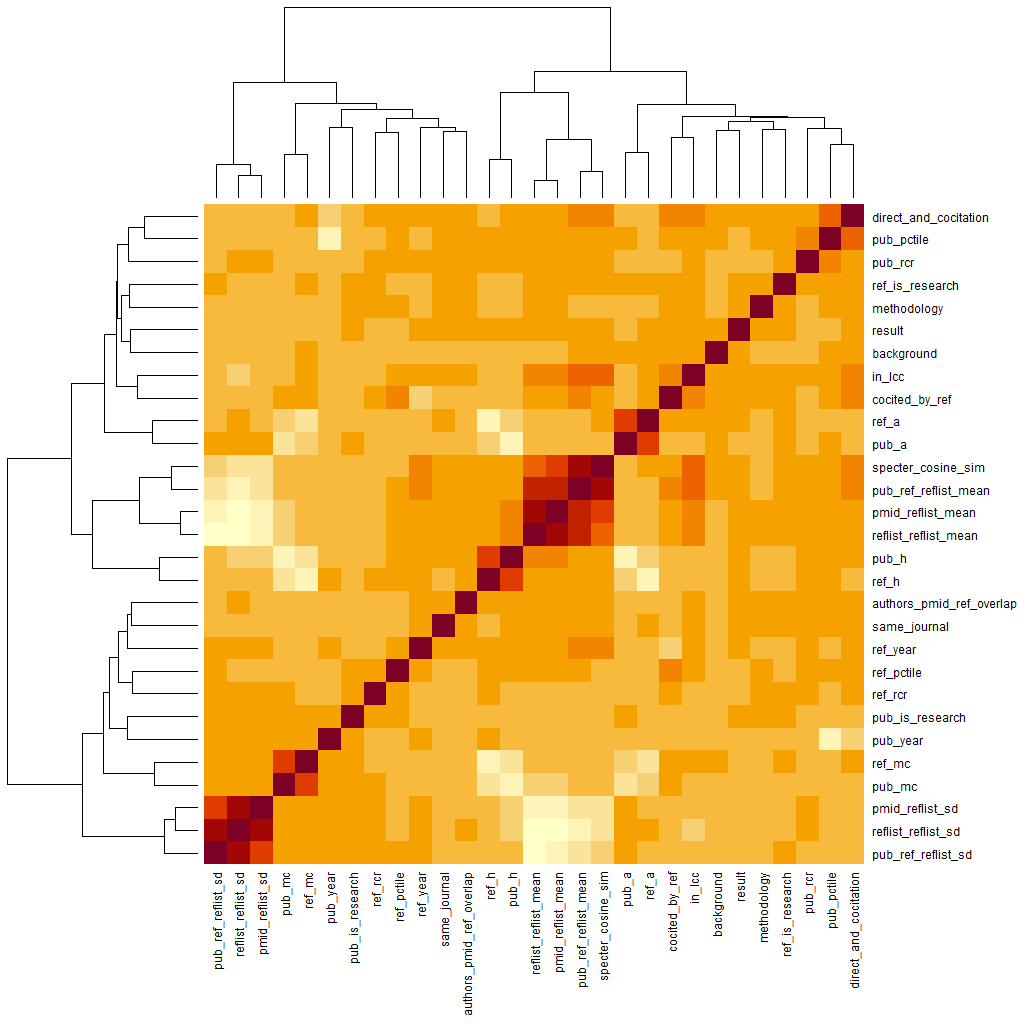

Supplemental Figure 1. Feature correlation matrix displayed as a heatmap, based on the training data.

#### Feature reduction analysis

To further test whether the model complexity could be reduced without sacrificing accuracy or external validity, we collapsed pub_year and ref_year into a composite variable dyear by subtracting ref_year from pub_year. Based on feature importance scores, we also removed two of the three local network structure features (direct_and_cocite and in_lcc). In addition, we removed the Human, Animal, and Molecular/Cellular scores, and retained only the percentiled RCR scores (pub_pctile and ref_pctile) while omitting the linear RCR scores. Finally, we removed the same_journal and is_research flags. We retained all the SPECTER data because these features received among the highest feature importance scores.

Our reduced feature set generated a model with similar levels of accuracy (F1 = 0.69 double check this). This indicates that raw accuracy was not much affected by the reduced feature space. However, when we checked its external validity on our five examples of known causal knowledge transfer (clinical drug progression, clinical disease progression, iPSC, XFP, and CRISPR), only four out of five examples showed the expected pattern of low prediction scores. This indicates that some external validity may have been traded off by reducing the feature set in this way. For this reason, we continued our experiments with the main model as described in the main text (full iCite + PubMed features but no Semantic Scholar features).
